# Genome-Wide Identification of Early-Firing Human Replication Origins by Optical Replication Mapping

**DOI:** 10.1101/214841

**Authors:** Kyle Klein, Weitao Wang, Tyler Borrman, Saki Chan, Denghong Zhang, Zhiping Weng, Alex Hastie, Chunlong Chen, David M. Gilbert, Nicholas Rhind

## Abstract

The timing of DNA replication is largely regulated by the location and timing of replication origin firing. Therefore, much effort has been invested in identifying and analyzing human replication origins. However, the heterogeneous nature of eukaryotic replication kinetics and the low efficiency of individual origins in metazoans has made mapping the location and timing of replication initiation in human cells difficult. We have mapped early-firing origins in HeLa cells using Optical Replication Mapping, a high-throughput single-molecule approach based on Bionano Genomics genomic mapping technology. The single-molecule nature and 290-fold coverage of our dataset allowed us to identify origins that fire with as little as 1% efficiency. We find sites of human replication initiation in early S phase are not confined to well-defined efficient replication origins, but are instead distributed across broad initiation zones consisting of many inefficient origins. These early-firing initiation zones co-localize with initiation zones inferred from Okazaki-fragment-mapping analysis and are enriched in ORC1 binding sites. Although most early-firing origins fire in early-replication regions of the genome, a significant number fire in late-replicating regions, suggesting that the major difference between origins in early and late replicating regions is their probability of firing in early S-phase, as opposed to qualitative differences in their firing-time distributions. This observation is consistent with stochastic models of origin timing regulation, which explain the regulation of replication timing in yeast.

## Introduction

The human genome is replicated from initiation events at on the order of 50,000 origins in each cell cycle (Masai et al., 2010). However, identifying the location and firing timing of human origins is an ongoing challenge (Hyrien, 2015). A major part of this challenge is the inefficient and heterogeneous nature of human origin firing (Berezney et al., 2000). In cases where mammalian origins are thought to occur at relatively well-defined sites, they appear to fire with as little as 10% efficiency (Dijkwel et al., 2002); in other cases, initiation appears to occur within broad initiation zones, spanning tens of kilobases (Hamlin et al., 2008). Therefore, population-based bulk approaches to mapping origins, which average origin-firing behavior across large populations of cells, must contend with low signal-to-noise ratios. As a result, although a number of bulk approaches have been used to map origins in the human genome (Cayrou et al., 2011; Besnard et al., 2012; Mesner et al., 2013; Foulk et al., 2015; Langley et al., 2016), there is little concordance between them (Mesner et al., 2013; Langley et al., 2016).

We have developed Optical Replication Mapping, a single-molecule technique to investigate spatial and temporal distribution of origin firing that combines the Bionano Genomics approach to mapping long individual DNA molecules (Lam et al., 2012) with *in vivo* fluorescent nucleotide pulse-labeling (Wilson et al., 2016) to directly visualize sites of replication initiation. This approach affords us excellent signal-to-noise characteristics, allowing us to identify origins that fire in a few as 1% of cells. We have used Optical Replication Mapping to identify and analyze early firing human replication origins.

## Results and Discussion

### Optical Replication Mapping of Early-Firing Human Origins

We mapped early-firing human replication origins by Optical Replication Mapping (ORM). We combined *in vivo* labeling of origins with Bionano visualization and mapping of long individual DNA molecules. Bionano analysis involves fluorescently labeling deproteinated DNA at the sites of the Nt.BspQI restriction enzyme, stretching individual molecules in nanochannels and imaging both the replication incorporation tracks and the restriction sites at about 1 kb resolution (Lam et al., 2012 and Figure 1A;). The pattern of fluorescent restriction sites allows the molecules and their associated replication tracks to be mapped to the human genome.

**Figure 1.**
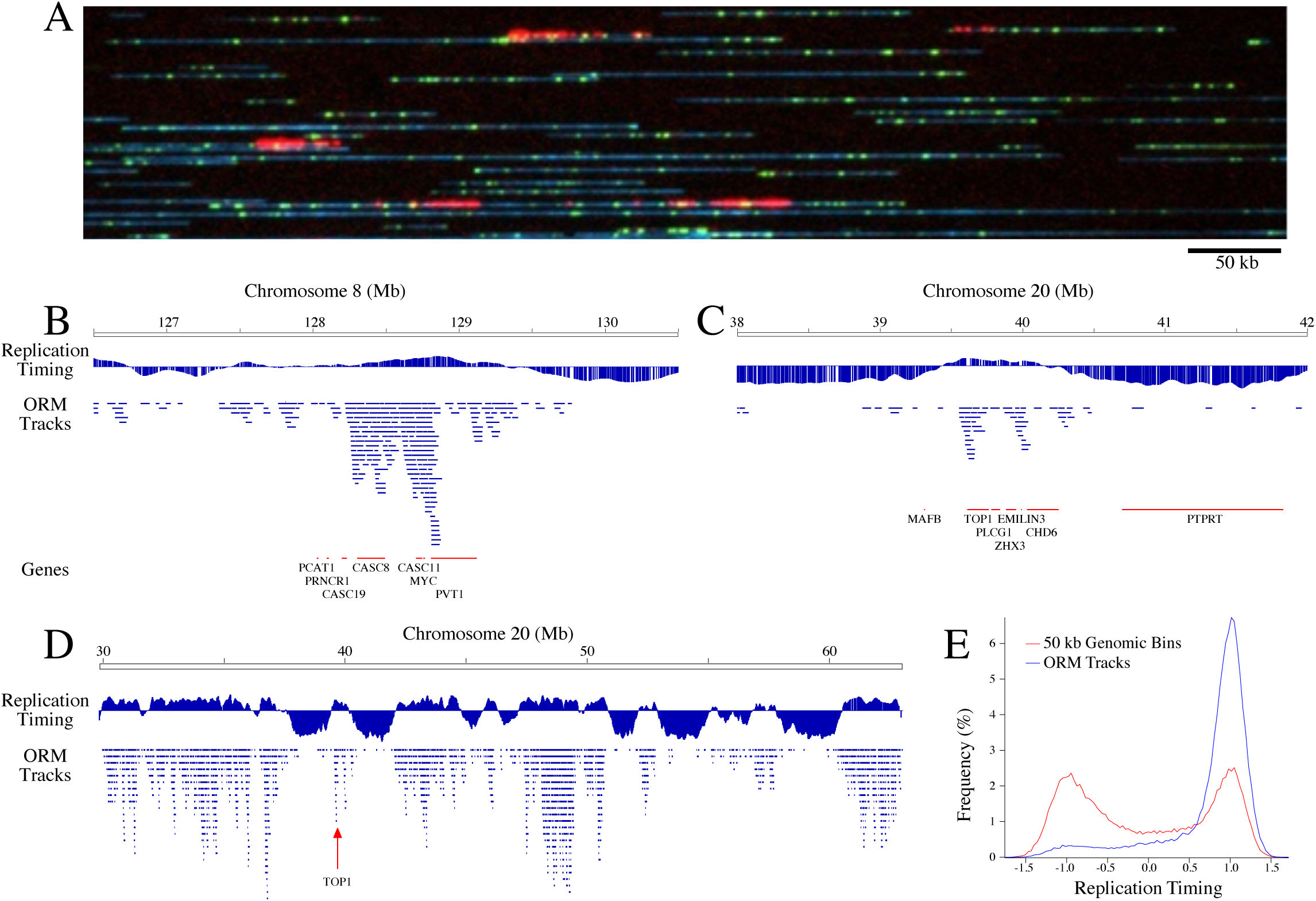
Optical Replication Mapping Identifies Early-Firing Origins. HeLa S3 cells were synchronized by mitotic shake off, arrested in early S-phase with aphidicolin, transfected with fluorescent dUTP and released from the aphidicolin arrest, allowing fluorescent nucleotide incorporation around early-firing replication origins. **A**) An image of the Bionano data with DNA in blue, Nt.BspQI restriction sites in green and incorporated nucleotides in red. The field of view is 650 kb. **B**,**C**) Distribution of replication tracks at two previously characterized human replication origins: Myc (B) and Top1(C). The distribution of replication tracks in shown beneath the HeLa S3 replication timing profile (Weddington et al., 2008) and above genes from the region. **D)** Distribution of replication tracks across the right arm of Chromosome 20, as in B. **E)** Replication timing of replication tracks compared to replication timing of each 50 kb bin in the genome.

HeLa S3 cells were synchronized in mitosis by mitotic shake off and arrested at the beginning of S phase with aphidicolin, which allows initiation at early-firing origins, but prevents elongation at those origins, and also prevents initiation at subsequent origins by activation of the intra-S-phase checkpoint (Feijoo et al., 2001). Cells were then transfected with Alexa647-dUTP, released from aphidicolin, allowed to recover overnight to complete replication and harvested for analysis on the Bionano Saphyr platform. Using this method, cells incorporate the transfected fluorescent nucleotides at origins that fired in the aphidicolin arrest and any that fire immediately after release. However, the transfected nucleotides are rapidly depleted, preventing incorporation at later-firing origins (Wilson et al., 2016). In this experiment, 87% of synchronized cells were in mitosis and 92% of them were fluorescently labeled, with 99% showing early replication patterns, indicating a high degree of synchrony and < 1% of contaminating asynchronous S phase cells.

We identified the fluorescent-nucleotide incorporation tracks and mapped them to the genome. The ORM tracks average 48.0±18.0 kb. Consistent with them reflecting sites of replication initiation, we find enrichment of signal around previously identified human replication origins (Figure 1B,C). In particular, we examined the distribution of replication tracks around the MYC and Top1 loci, both characterized as early-firing origins in the human genome (Tao et al., 2000; Keller et al., 2002), and found a pronounced enrichment at these loci, particularly in the intergenic regions surrounding these genes. Furthermore, our signals are enriched in the earliest replicating parts of the genome, as defined by replication timing profiling (Weddington et al., 2008), as expected for early-firing origins (Figure 1D,E).

### Genome-Wide Mapping of Early-Firing Human Origins

To map the genome-wide distribution of early-firing human replication origins, we collected 290x coverage of early-firing origin mapping data. We identified 79267 incorporation tracks in this dataset, for an average origin firing density of about 270 early-firing origins per cell. However, across the dataset 90% of the 6166 early-firing initiation zones identified by OK-seq (Petryk et al., 2016) were detected by ORM tracks, demonstrating that, in the population, most or all early-firing origins fire in the aphidicolin arrest. This result shows that early-origin firing is quite heterogeneous, with only about 4% (270/6166) of early-firing origins firing in any given cell before origin firing is inhibited by the aphidicolin-induced checkpoint.

A major advantage of optical replication mapping is that it records the number of fibers that both have and do not have replication tracks, allowing for the calculation of the absolute efficiency with which replication initiates at each locus in the genome. To investigate the frequency of early origin firing across the genome, we examined the frequency of replication initiation, as measured by optical replication mapping data, at all 9830 initiation zones (both early and late firing) previously identified by Okazaki-fragment mapping (OK-seq) (Petryk et al., 2016). We find that the local efficiency of firing in these zones ranges up to a maximum of approximately 5%, with the majority of the more efficient zones found in early-replicating regions of the genome, as determined by replication timing profiling (Weddington et al., 2008), consistent with the results in Figure 1E (Figure 2A). Although this number is low, it is unlikely that it is a severe under estimate. It seems unlikely that we are under counting replication tracks because, assuming the all forks incorporate the fluorescent nucleotide at the same rate, all tracks should have similar signal strength. Consistent with this assumption, we see a roughly normal distribution of track lengths and signal intensities (data not shown). Likewise it seems unlikely that we are over counting fibers without replication tracks. Even if the 13% of our cells that were not synchronized in mitosis and the 8% that did not incorporate nucleotide all contributed to over counting of fibers without replication tracks, the maximum initiation rate would only increase to about 6%. More likely, the aphidicolin simply arrests cell quickly, before many origins have a chance to fire.

**Figure 2.**
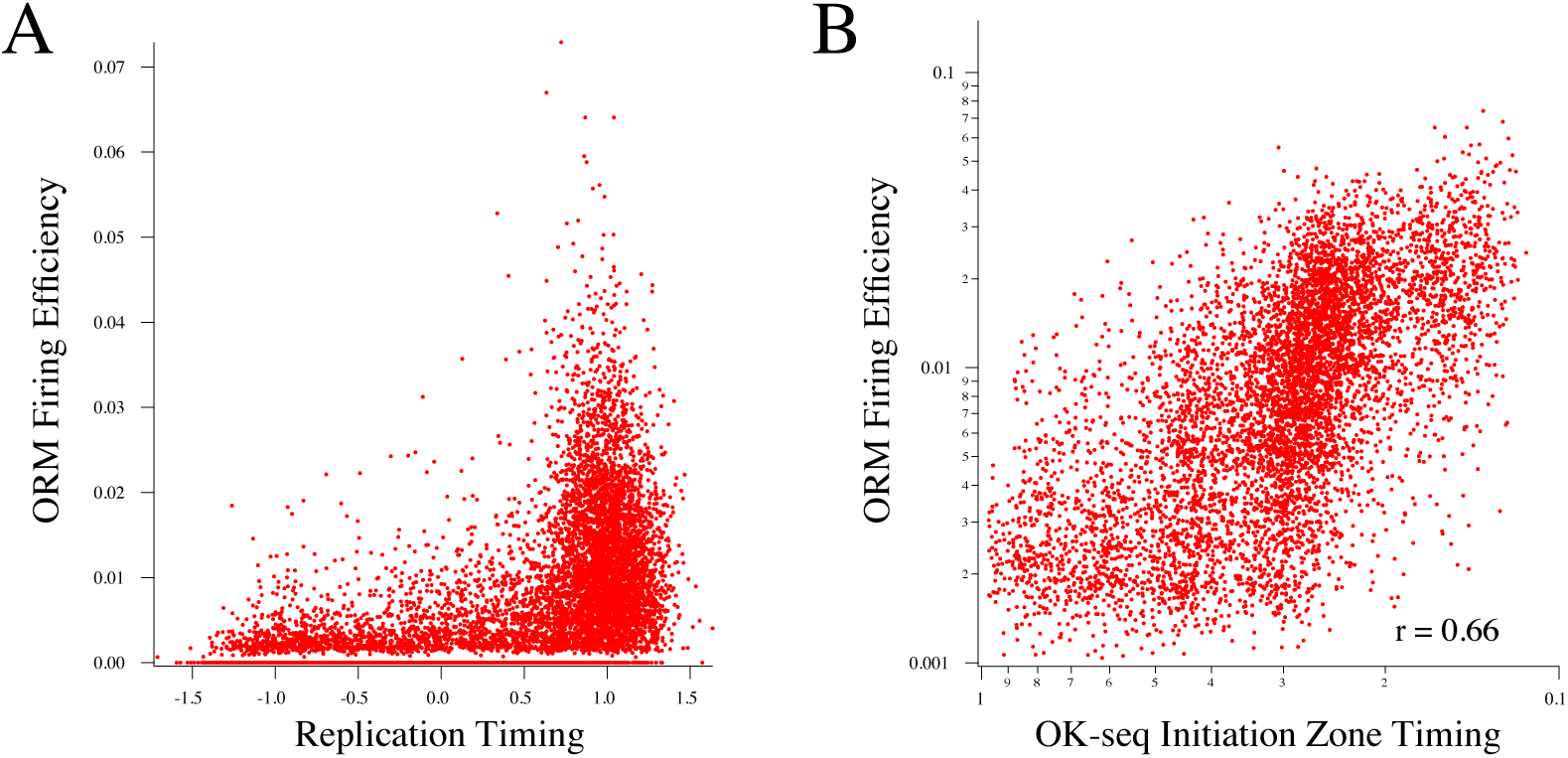
Replication Initiation Efficiency of Initiation Zones. **A)** 9830 initiation zones identified by OK-seq analysis (Petryk et al., 2016) are plotted as replication timing (Weddington et al., 2008) versus firing efficiency, calculated as the number of replication tracks overlapping each initiation zone divided by the number of fibers covering each zone. **B)** The firing efficiency of the same 9830 zones are plotted versus the initiation time of each zone, as determined by OK-seq analysis (Petryk et al., 2016). Initiation timing differs from replication timing in that replication timing includes the effect of passive replication by forks initiating outside of the zone, while initiation timing measures only initiation from within the zone.

To directly test if the firing efficiently we measure by optical replication mapping is correlated with the initiation times of the OK-seq initiation zones, we compared firing efficiency with the initiation time of each initiation zone, as determined by OK-seq (Petryk et al., 2016). Initiation timing differs from replication timing in that replication timing includes the effect of passive replication by forks initiating outside of the zone, while initiation timing measures only initiation from within the zone. We find a significant correlation between the two (Figure 2B, r = 0.66), consistent with early firing origins having a higher efficiency of firing in early S phase.

Visual inspection of the distribution of ORM tracks suggests that initiation in human cells is not concentrated at discrete, frequently used origins. Instead initiation appears to be distributed across broad initiation zones, as has been observed at specific loci and in other genome-wide analyses (Dijkwel et al., 2002; Hamlin et al., 2008; Besnard et al., 2012; Mesner et al., 2013; Petryk et al., 2016). Even the initiation zone at the Myc locus, which fires in 4.9% of cells, appears to have initiation events distributed over tens of kilobases (Figure 1B). However, we have not yet performed a rigorous analysis to test this contention.

### Comparison of Different Origin Mapping Datasets

We compared our optical replication mapping data to four published HeLa replication-initiation-mapping datasets: OK-seq (Petryk et al., 2016), SNS-seq (Picard et al., 2014), Ini-seq (Langley et al., 2016) and Orc1 ChIP-seq (Dellino et al., 2013). By visual inspection, our ORM tracks appear to be enriched around the initiation zones identified by OK-seq (Figure 3A). OK-seq measures the ratio of replication fork directions (RFD) at every point in the genome (Petryk et al., 2016). Transitions from leftward-moving forks to rightward-moving forks identify zones of initiation. Across the genome, we see colocalization of OK-seq initiation zones and ORM tracks. We see similar colocalization with Ini-seq signal, a cell-free initiation mapping approach, and Orc1 ChIP-seq, and to a lesser extent, colocalization with SNS-seq, which maps initiation events by sequencing short nascent strands produced by replication initiation (Figure 3A).

**Figure 3.**
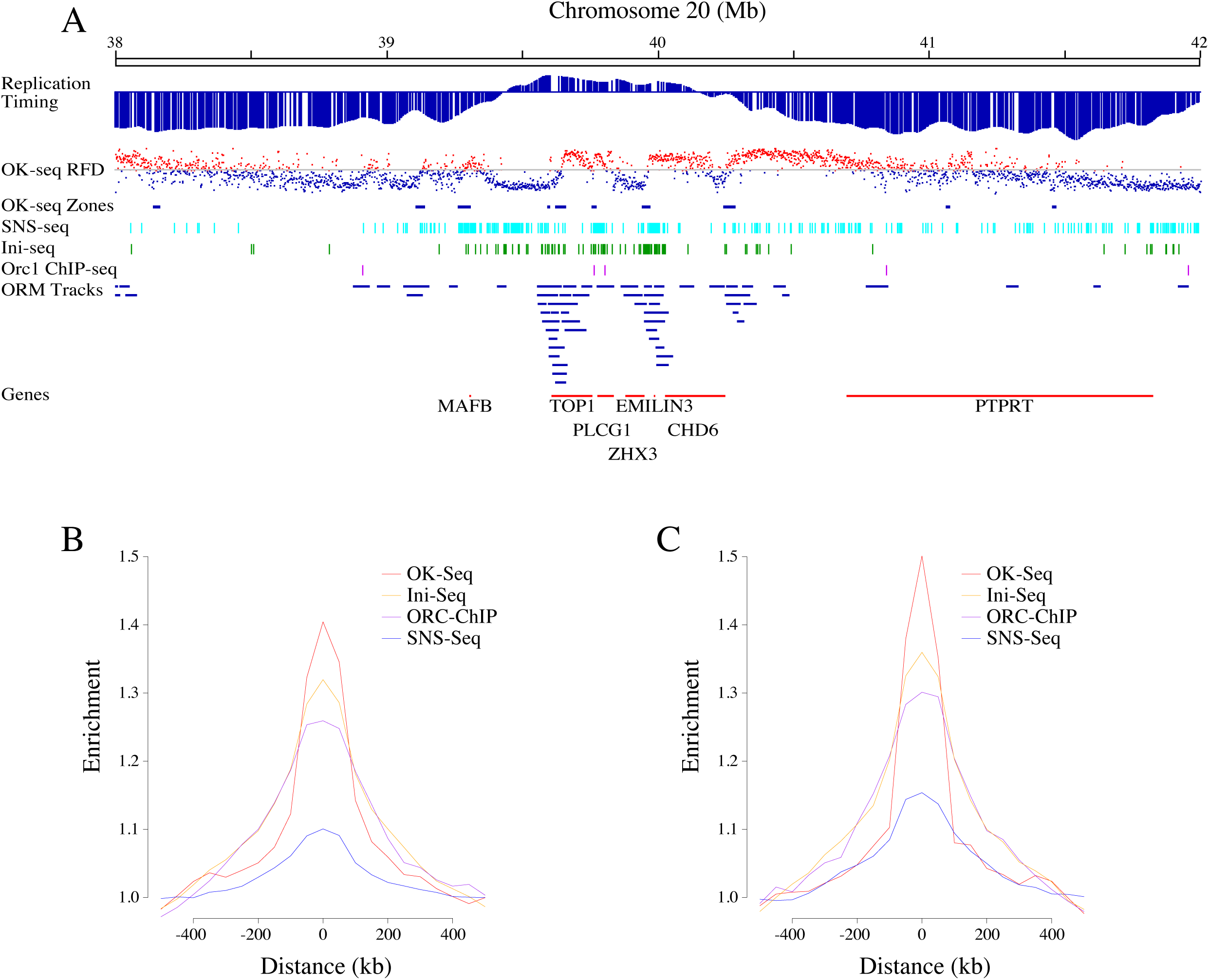
Comparison of Optical Replication Mapping with Other Replication Initiation
Mapping Datasets. **A)** Comparison of optical replication mapping tracks, OK-seq replication fork directionality (RFD) showing points with predominantly leftward-(blue) or rightward-(red) moving forks (Petryk et al., 2016), OK-seq initiation zones calculated from the RFD data as zones of transition from leftward-to rightward-moving forks (Petryk et al., 2016), SNS-seq (Picard et al., 2014), Ini-seq (Langley et al., 2016) and Orc1 ChIP-seq (Dellino et al., 2013) at the Top1 locus. **B)** Enrichment of optical replication mapping tracks at relative to the other initiation mapping datatypes. **C)** Enrichment of the other initiation mapping datatypes relative to optical replication mapping tracks. The slightly higher enrichment of other datatypes around the optical replication mapping tracks suggests that the optical replication mapping dataset has fewer false-positive signals.

To determine if the apparent co-localization of initiation mapping data is robust, and to quantitate its extent, we calculated the enrichment of ORM tracks around the other replication initiation signals and, conversely, the enrichment of those initiation signals around ORM tracks. We find that ORM tracks colocalize most extensively with OK-seq initiation zones, to a similar extent with Ini-seq and Ocr1 ChIP-seq signal and to a lesser extent with SNS-seq signal (Figures 3B,C). The slightly higher enrichment of other datatypes around the optical replication mapping tracks (Figure 3C) compared with enrichment of the optical replication mapping tracks around the other datatypes (Figure 3B) suggest that the optical replication mapping dataset has fewer false-positive signals than the other datasets.

### Early-firing of Origins in Late-Replicating Regions

Although 90% of our early-firing origins fire in early replicating regions of the genome, 10% fire in late-replicating regions (Figure 1E). By visual inspection, the ORM tracks in late-replicating parts of the genome appear to be in the locally earliest-replicating parts of the late-replicating regions (for instance at 127 Mb in Figure 1B and 39.1 Mb in Figure 1C), suggesting that they reflect the location of the origins normally used to replicate these regions. To test this enrichment, we divided the genome into early and late replicating regions, normalized to the distribution of replication timing and re-calculated the enrichment of ORM tracks in the relatively early-and late-replicating regions of both parts of the genome. We find that ORM tracks significantly enriched in the locally earliest-replicating regions of the late-replicating part of genome (Figure 4A), consistent with these early-firing events in late-replicating region being rare examples of normally late-firing origins firing early. Moreover, we find the ORM tracks are depleted in the later-replicating regions of the early-replicating part of the genome, which are largely passively replicated.

**Figure 4.**
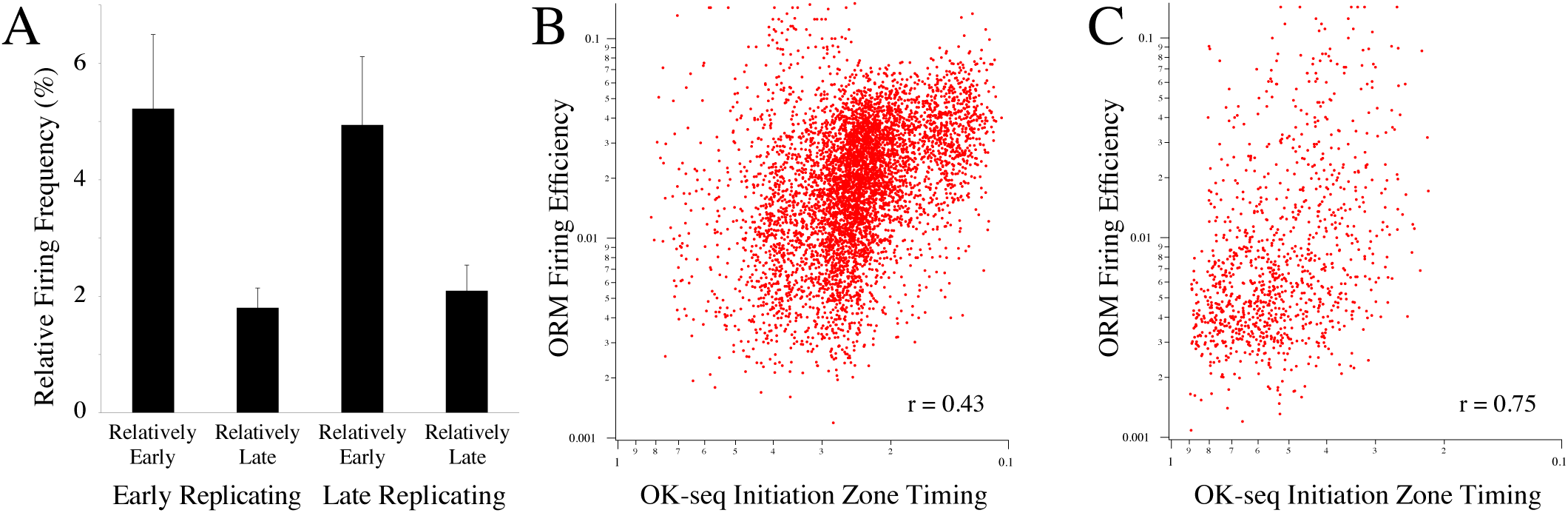
Distribution of Early-Firing Replication Tracks in Late-Replicating Regions of the Genome. **A**) The relative frequency of firing in the relatively early-and late-replicating regions of the early-and late-replicating halves of the genome (see Methods for details of the calculation). **B**,**C**) The firing efficiency of OK-seq initiation zones are plotted versus the initiation times of each zone, as in Figure 2B, for zones in early-(B) and late-(C) replicating regions of the genome, as determined by replication profiling (Weddington et al., 2008).

As an alternative analysis, we investigated whether the correlation between initiation zone timing and ORM track frequency seen in Figure 2B holds for late-replicating regions of the genome. We calculated the correlation between the timing of initiation zone activation calculated from OK-seq data and the frequency with which ORM tracks are observed at those initiation zones separately for early-and late-replicating regions of the genome. We find a strong correlation between initiation zone timing and ORM track frequency in both regions (Figure 4B,C), suggesting that when origins in late-replicating regions fire early in S phase, they do so at the sites of the most efficient late-firing origins.

Such a distribution of early-firing origins—frequent firing at early-replicating origins and rare firing at late-replicating origins—is reminiscent of the stochastic regulation of origin-firing timing seen in budding yeast (Czajkowsky et al., 2008; de Moura et al., 2010; Yang et al., 2010). Our results are consistent with simple stochastic models of replication timing in which the only difference between early-and late-firing origins is their probability of firing. More efficient origins are more likely to fire first and thus, on average, have earlier firing times, but sometimes they still fire late, whereas less efficient origins are less likely to fire early and thus, on average, have later firing times, but sometimes they can fire early (Rhind et al., 2010). Further analysis of our optical replication mapping data and future optical replication mapping experiments will shed additional light on this question.

## Methods

### Cell Labeling and Sample Preparation

HeLa S3 cells were grown in DMEM plus 10% Cosmic Calf Serum (GE Life Science SH30087) and Pen/Strep. Cells were synchronized at metaphase by incubation with 0.05 µg/ml nocodazole for 4 hours followed by shake off. Metaphase cells were released into fresh media containing 10 µg/ml aphidicolin (Millipore Sigma 178273) for 16 hours. Cells were then washed with PBS, trypsinized, and nucleofected (Lonza, kit SE, HeLa S3 HV program) in the presence of Aminoallyl-dUTP-ATTO-647N (Jena Bioscience NU-803-647N). Cells were recovered in incubator overnight to allow nucleotide incorporation. Cells were detached with trypsin and snap frozen in liquid nitrogen.

### Optical Replication Mapping

Cell pellets were thawed quickly at 37ºC, resuspended in Cell Buffer (Bionano 20340), and embedded into low-melting point agarose plugs using the CHEF Mammalian Genomic DNA Plug Kit (Bio-Rad #1703591) according to the manufacturer’s instructions, with a target concentration of 300,000 cells per plug. Cells immobilized in plugs were lysed and treated with Proteinase K and RNase A. The plugs were then melted and solubilized, and the resulting DNA was further cleaned by drop dialysis. The DNA was fluorescently labeled with green fluorophores at Nt.BspQI sites and counterstained with YOYO-1 using Bionano Genomics’ Nicks, Labels, Repairs and Stains (NLRS) labeling kit (Bionano RE-012-10). Throughput the entire process, the samples were shielded from light whenever possible, in order to maintain integrity of the fluorophores.

Labeled DNA samples were loaded onto an Irys chip (Bionano FC-020-01) or Saphyr chip (Bionano Genomics FC-030-01) and run on the corresponding Irys or Saphyr instrument (Bionano Genomics IN-011-01), following the manufacturer’s instructions. Briefly, the DNA was coaxed into an array of parallel nanochannels via electrophoresis, thereby elongating the DNA to a uniform contour length, allowing for accurate measurement of distances along the molecules. Molecules were imaged to collect the YOYO-1 DNA signal in a false-color blue channel, the Nt.BspQI label in the green channel, and the *in vivo* replication incorporation tracks in the red channel.

Images were converted to digitized molecules and the positions of the green and red labels on each fiber were determined using Bionano Genomics data analysis pipeline. The Nt.BspQI site labels were aligned to human reference genome hg19 by creating an *in silico* digestion of the reference FASTA. Alignment of the fibers' Nt.BspQI sites allowed mapping of the red replications signal to the genome. The pipeline produces one .bnx file containing the intensity and location of each green and red signal on each fiber and one .xmap file containing the genomic location of each fiber. The Bionano data is available on line <https://www.dropbox.com/sh/kndhowsvde1qeyo/AABRunjknMUwVKAPZZ7Y5ZEXa?dl=0>.

### Identification of Initiation Sites

To identify replication tracks, we used custom scripts <https://github.com/tborrman/DNA-rep/tree/master/BioNano> to filter out spurious red signals and segment the remaining red signals into identified replication tracks. In brief, direction_bed_seg_new_format.py takes the .bnx and .xmap files as input and outputs three files: (1) output.bed, in which each row corresponds to an identified replication track and includes the molecule ID, track polarity, polarity strength, and sum of red label signal in the track. Background red signals were removed using the following criteria. We filtered for molecules containing a red label with at least five neighboring red labels within 20kb. We then kept only the signals from these five-neighbor red labels and their associated neighbors to incorporate in our filtered segments. Segments were then split if distance between one red label and next label was greater than or equal to 30kb. (2) unfiltered.bedGraph for all the red label signals. (3) filtered.bedGraph for the filtered red label signals. Further information on output file formats and specifications can be found in the README.md <https://github.com/tborrman/DNA-rep/blob/master/README.md>. For faster processing direction_bed_multi_controller.py was used as a controller for running direction_bed_seg_new_format.py in parallel on an LSF cluster environment. For generation of signal tracks and bam segment files for figures we used the script cleanbed_for_bed.py along with a set of commands from bedtools v2.17.0 and samtools v0.1.19. These commands are also specified in the above mentioned README.md. Filtered replication signals and mapped replication tracks were visualized in IGV (Robinson et al., 2011) and are available online <https://www.dropbox.com/sh/kndhowsvde1qeyo/AABRunjknMUwVKAPZZ7Y5ZEXa?dl=0>.

### Comparative Analysis of Origin Localization

Published genome-wide HeLa cell replication-initiation-mapping datasets—OK-seq (Petryk et al., 2016), SNS-seq (Picard et al., 2014), Ini-seq (Langley et al., 2016) and Orc1 ChIP-seq (Dellino et al., 2013)—were used to compute enrichments around the ORM tracks and surrounding regions. When necessary, genomic coordinates were remapped to hg19 using LiftOver. The initiation times of the OK-seq initiation zones were computed as indicated (Petryk et al., 2016). Replication timing data was obtained from ReplicationDomain.org. Features were visualized in IGV (Robinson et al., 2011).

### Calculation of Relative Firing Frequency in Early and Late-Replicating Regions

The genome was divided into early-replicating (0 to 1) and late-replicating (-1 to 0) by replication timing (Weddington et al., 2008). Each region was subdivided into 28 timing windows. The number of replication tracks in each window was normalized to the number of 50 kb genomic bins in each window. The probability density function of replication tracks in each window of the two regions was calculated and the average and standard deviation of windows in the early and late half of each region was plotted in Figure 4A.

## Author Contributions

**Conceptualization:** DG, NR - **Methodology:** KK, WW, TB, ZW, AH, CC, DG, NR - **Investigation:** KK, WW, TB, SC, DZ - **Software:** WW, TB, ZW, CC - **Formal Analysis:** WW, TB, CC, NR - **Resources:** ZW, AH, CC, DG, NR - **Data Curation:** WW, TB, CC - **Writing – Original Draft:** NR - **Writing – Review & Editing:** KK, WW, TB, SC, DZ, ZW, AH, CC, DG, NR - **Visualization:** WW, CC, NR - **Supervision:** ZW, AH, CC, DG, NR - **Project Administration:** NR - **Funding Acquisition:** ZW, AH, CC, DG, NR

